# Lysosomal calcium loading promotes arrhythmias by potentiating ryanodine receptor release

**DOI:** 10.1101/2022.11.09.515777

**Authors:** Zhaozheng Meng, Rebecca A Capel, Samuel J Bose, Erik Bosch, Sophia de Jong, Robert Planque, Antony Galione, Rebecca AB Burton, Alfonso Bueno-Orovio

## Abstract

Spontaneous calcium release by ryanodine receptors (RyRs) due to intracellular calcium overload results in delayed afterdepolarisations, closely associated with life-threatening arrhythmias. In this regard, inhibiting lysosomal calcium release by two-pore channel 2 (TPC2) knockout has been shown to reduce the incidence of ventricular arrhythmias under β-adrenergic stimulation. However, mechanistic investigations into the role of lysosomal function on RyR spontaneous release remain missing. We investigate the calcium handling mechanisms by which lysosome function modulates RyR spontaneous release, and determine how lysosomes are able to mediate arrhythmias by its influence on calcium loading. Mechanistic studies were conducted using a population of biophysically-detailed mouse ventricular models including for the first time modelling of lysosomal function, and calibrated by experimental calcium transients modulated by TPC2. We demonstrate that lysosomal calcium uptake and release can synergistically provide a buffering pathway of fast calcium transport, by which lysosomal calcium release primarily modulates sarcoplasmic reticulum (SR) calcium reuptake and RyR release. Enhancement of this lysosomal transport pathway promoted RyR spontaneous release by elevating the SR-junction calcium gradient. In contrast, blocking either lysosomal calcium uptake or release revealed an antiarrhythmic impact. Under conditions of calcium overload, our results indicate that these responses are strongly modulated by intercellular variability in L-type calcium current, RyR release, and SERCA reuptake. Altogether, our investigations identify that lysosomal calcium handling directly influences RyR spontaneous release by regulating the SR-junction calcium gradient, suggesting antiarrhythmic strategies and identifying key modulators of lysosomal proarrhythmic action.

**Statement of Significance:** Delayed afterdepolarisations arising from spontaneous RyR calcium release are an important risk factor for arrhythmogenesis. Inhibiting lysosomal calcium release by TPC2-KO reduces the propensity for ventricular arrhythmias. However, understanding downstream effects of lysosomal calcium release on spontaneous RyR release is lacking. Understanding lysosomes as arrhythmia sources requires alternative approaches to controlled laboratory techniques: these restrain variability experimentally and statistically. Our study presents two methodological novelties by focusing on calibration with experimental findings using a population of biophysically-detailed models and incorporating lysosomal mechanisms. Lysosomal calcium handling promotes RyR spontaneous release by elevating the SR-junction calcium gradient. Blocking lysosomal function uncovers an antiarrhythmic strategy. Lysosome-release proarrhythmic risk is determined by synergistic enhancements of lysosomal uptake with RyR release or L-type calcium current.

## Introduction

Disruption in calcium signalling underlies many cardiac disorders, including contractile dysfunction and rhythm disorders [1-4]. In cardiomyocytes, the process of excitation-contraction coupling links depolarisation of the plasma membrane to muscle fibre contraction [5]. Membrane depolarisation in response to the arrival of an action potential leads to the activation of voltage gated L-type calcium channels located within the sarcolemma and transverse (t-)tubules. The resulting inward calcium current (I_CaL_), causes an increase in calcium concentration within the subsarcolemmal junctional space, located between the surface membrane and the sarcoplasmic reticulum (SR). This elevation in junctional calcium activates ryanodine receptors (RyR) which then release calcium from the SR into the cytosol. If sufficient RyR are activated, this process of calcium-induced calcium release (CICR) leads to a cellular calcium transient characterised by an initial rise in cytosolic calcium followed by a fall resulting from the combination of RyR closure, reuptake of calcium into SR via the SR calcium-ATPase (SERCA), and release from the cell via the sodium-calcium exchanger (NCX). Calcium transients are linked to cell contraction due to the binding and activation of cytosolic calcium to troponin on the muscle filaments [5]. Tight regulation of cellular calcium is therefore essential in cardiac cells in order to maintain healthy rhythm and contractility.

Calcium in cardiomyocytes is primarily buffered within the SR. However, other organelles including lysosomes and mitochondria may also act as calcium stores and influence calcium handling [6-8]. In addition to β-adrenergic signalling, calcium signalling undergoes regulation via multiple alternative pathways and messengers, including nicotinic acid adenine dinucleotide phosphate (NAADP), inositol-1,4,5 trisphosphate, and cyclic ADP ribose [3, 6-9].

The involvement of lysosomes in cardiac disorders has long been recognised. In the 1960s, higher numbers of lysosomes were observed in the atrial tissue of chronically diseased hearts from human patients [10], and lysosomes were increased in canine atria following introduction of atrial septal defects [11]. Multiple studies have since demonstrated that lysosomal dysfunction can be linked to cardiac disorders including arrhythmia and heart failure [9, 12-14]. Lysosomal calcium release occurs primarily through the opening of type-2 two-pore channels (TPC2) following activation by NAADP [13, 15-18]. Transmission electron microscopy has shown that in cardiomyocytes, lysosomes are located within close proximity to the SR, mitochondria and t-tubules, and may form signalling microdomains involving membrane contact sites with these organelles [19]. Release of lysosomal calcium via TPC2 in response to NAADP enhances RyR calcium release from the SR [20], and inhibition of the NAADP pathway has been shown to be protective against β-adrenergic induced arrhythmias [21]. Furthermore, lysosomes have been shown to generate oscillating calcium currents that can trigger potentially lethal cardiac arrhythmias, and these oscillations can also be supressed by the inhibition of NAADP [22]. Despite these observations, the role of lysosomal calcium signalling in cardiomyocytes has been relatively understudied and lysosomes are not included in current mathematical models of cardiac calcium handling [23, 24]. This severely constrains our understanding of their possible proarrhythmic role and our capability to suggest appropriate antiarrhythmic therapies.

Whilst separate components of calcium signalling within cardiomyocytes can be investigated experimentally in isolation, mathematical modelling enables comprehensive predictive studies to be performed consistently by integrating these components [23, 24]. Models of cardiac excitability have several benefits compared to physiological experiments. Biological experiments are subject to the combined limitations of loss of variability, for example through the averaging of data, as well as limited sample size and throughput [25-27]. Mathematical models based purely on experimental data can therefore of course also suffer from the loss of individuality resulting from the averaging of biological data [25]. However, the use of more recent advances in approaches for studying variability such as experimentally calibrated populations-of-models frameworks, enables such biological variability to be incorporated into models of cardiac electrophysiological signalling [26, 27]. Here we modify an existing model of mouse ventricular calcium handling by Morotti *et al*. [24] to include the lysosomal compartment based on experimental data on spatial organisation and lysosomal action. In our updated model we adopt a population of models-based approach to account for variability in electrophysiology and calcium handling [26, 27]. This model was used to investigate mechanisms of lysosomal calcium handling in arrhythmogenesis and RyR spontaneous release under conditions of calcium overload (hypercalcemia and fast pacing) in conjunction with β-adrenergic stimulation. Our modelling data supports the hypothesis that lysosomes can act as an additional intracellular calcium store, providing an additional buffer for intracellular calcium, and can influence the driving force for spontaneous calcium release from the SR to the junctional space. Identifying new targets in arrhythmic processes is therefore of great importance and has potentially high clinical value.

## Materials and Methods

### Mathematical modelling and simulation studies

The model by Morotti *et al*. [24] was used as the baseline mathematical model for our investigations because of matching species and the inclusion of β-adrenergic response and CaMKII signalling. A lysosomal calcium compartment was formulated into the model to account for lysosomal calcium handling, as detailed next. The simulated results matched calcium transients and action potentials in the experimental results [28]. Each simulation was run for 150 beats at 1 Hz pacing rate to allow the system to enter steady-state, and all reported biomarkers were calculated on the last beat, unless otherwise specified. NAADP concentration (*[NAADP]*) was used as input parameter to regulate TPC2 release, with values of 1 nM for control (CTRL) and 15 nM to simulate nearly fully-open probability under either exposure to isoprenaline (ISO) or NAADP-AM (cell permeant analogue of NAADP). An ISO ligand value of 100 nM was used in the simulated ISO protocols, which matched the predefined ISO responses of the Morotti *et al*. model with the magnitude of β-adrenergic response observed in our experimental data.

### Modified ventricular cell model with lysosome compartment

Figure 1 illustrates the integration of the new lysosomal compartment into the Morotti *et al*. model. Based on organelle localisation and lysosome action in experimental findings [18, 19, 28, 29], the lysosomal compartment was included in the model as an additional calcium pool, incorporating calcium diffusion with the neighbouring junctional and cytosolic subspaces, as well as a calcium loading channel between the junctional and lysosomal subspaces, and lysosomal calcium release through TPC2 into the cytosolic subspace.

**Figure 1.**
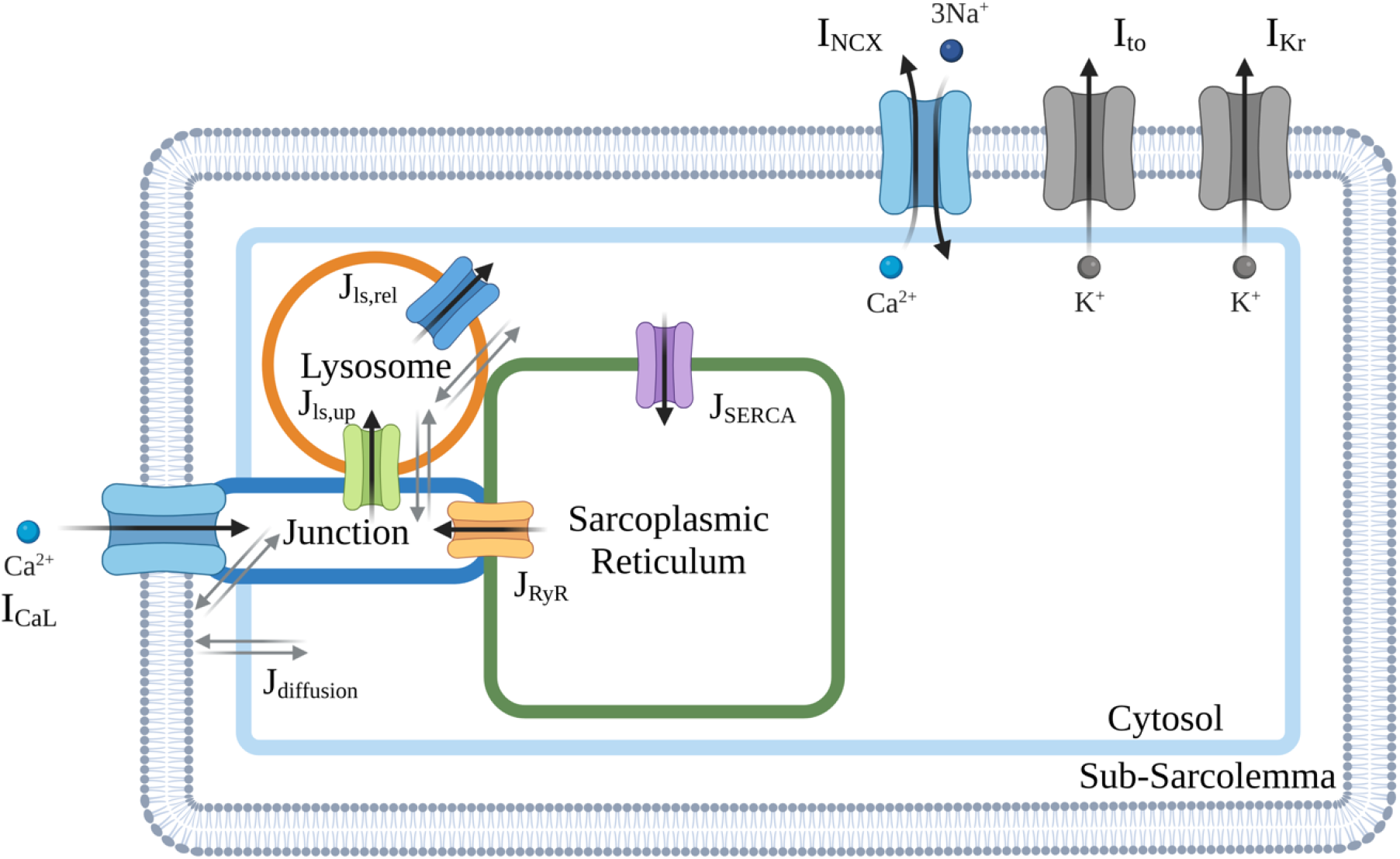
Schematic diagram of modified ventricular cell model with lysosome compartment. Cellular electrophysiology of mouse ventricular cardiomyocyte is represented in the existing Morotti *et al*. [24] model with β-adrenergic stimulation and CaMKII signalling. Only the main ion channels varied in a population of models approach in this study are shown. A more complete overview of the model can be found in Morotti *et al*. [24].

The lysosomal calcium inward flux was modelled as a calcium loading channel (CLC) between the junctional and lysosome subspaces. The lysosomal calcium release through TPC2 was modelled as a lysosomal calcium efflux from the lysosome compartment into the cytosolic subspace. These are respectively given by:

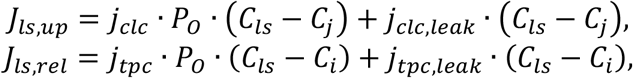

where *j*_*clc*_ and *j*_*tpc*_ denote the maximal CLC uptake and TPC2 release flux densities into the lysosomal and cytosolic subspaces; *C*_*ls*_, *C*_*j*_, and *C*_*i*_ are the calcium concentrations in the lysosomal, junctional, and cytosolic subspaces, respectively; *j*_*clc,leak*_ and *j*_*tpc,leak*_ denote the leak rates of calcium through the CLC and TPC2 channels to the junctional and cytosolic subspaces, respectively; and *P*_*O*_ represents the open probability of the lysosomal uptake and release channels. Based on previous experimental data [18, 28], we adopt the formulation of Penny *et al*. [30] to describe the open probably of TPC2 channels. This is given by:

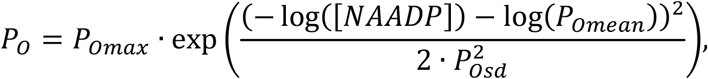

so that *P*_*O*_ follows a Gaussian distribution on the NAADP concentration, *P*_*Omax*_ denotes its maximum open probability, and *P*_*Omean*_ and *P*_*Osd*_ represent the mean and standard deviation of the distribution. For CLC channels, given the scarcity of experimental datasets, we adopted an analogous formulation to TPC2 channels. Such an assumption correctly recapitulated our calcium transient data under CTRL, NAADP-AM and ISO protocols (see Figure 2). This modelling assumption can nevertheless be easily revisited, shall additional datasets on lysosomal calcium loading become available.

**Figure 2.**
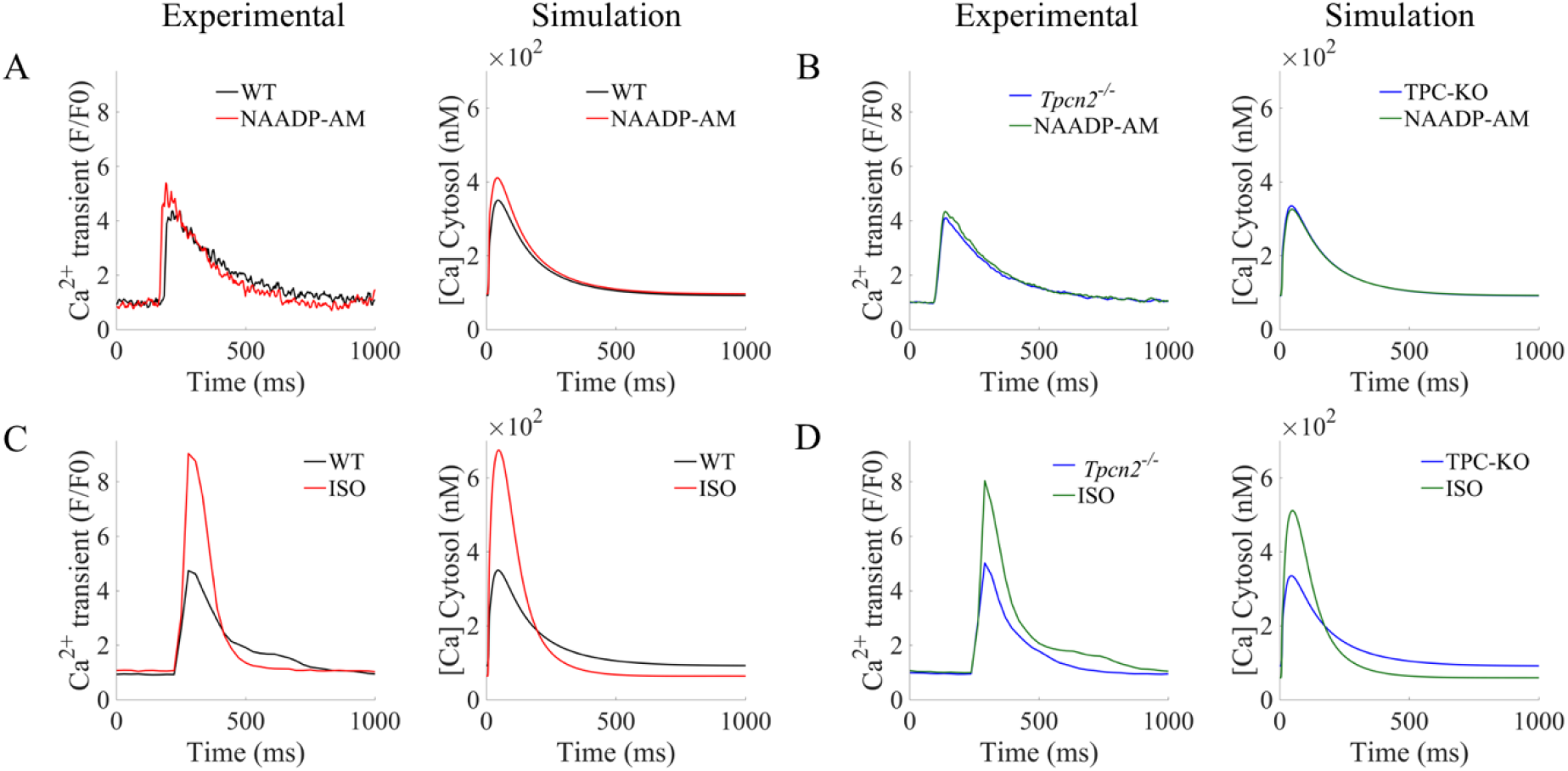
Validation of lysosomal calcium handling model against pharmacological protocols. **2A, 2B:** Calibration of Ca^2+^ transients under increased lysosomal release by NAADP-AM (WT vs TPC2-KO, respectively). **2C, 2D:** Validation of Ca^2+^ transients under β-adrenergic stimulation by ISO (WT vs TPC2-KO, respectively).

The inclusion of the new lysosomal calcium fluxes then yields the following system of ordinary differential equations for conservation of mass of calcium ions:

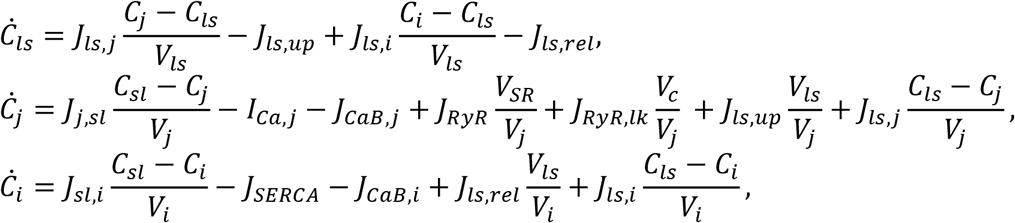

where *Ċ*_*ls*_, *Ċ*_*j*_, and *Ċ*_*i*_ denote the time derivatives of calcium concentrations in the lysosomal, junctional, and cytosolic subspaces, respectively, while *V*_*ls*_, *V*_*j*_, *V*_*SR*_, and *V*_*i*_ represent the volumes of the lysosomal, junctional, SR, and cytosolic subspaces. In the equations above, *I*_*Ca,j*_ denotes the junctional calcium fluxes via calcium currents, *J*_*CaB,j*_ are the junctional calcium buffers, *J*_*RyR*_ and *J*_*RyR,lk*_ the release and leak fluxes through ryanodine receptors, respectively, *J*_*SERCA*_ denotes the calcium reuptake fluxes by SERCA, and *J*_*CaB,i*_ the cytosolic calcium buffers. In these equations, *J*_*ls,j*_ and *J*_*ls,i*_ are the rates of diffusion fluxes between the lysosome calcium pool and junctional or cytosolic subspaces, respectively.

The remaining equations in the Morotti *et al*. model [24] remained unaltered, including those describing calcium concentrations in the SR and sarcolemmal compartments. Parametrisation of the lysosomal compartment is provided in Supplementary Material Table S1.

### Experimental data

The experimental datasets include calcium transients and action potentials recordings at 1 Hz under CTRL, NAADP-AM and ISO protocols in both WT and TPC2-KO mouse ventricular cardiomyocytes, and have been reproduced with permission from the original authors [28, 31].

### Cellular variability in electrophysiology and calcium handling

We adopted a population of models approach [26, 27] to account for cellular variability in electrophysiology and calcium handling. Such an approach has been especially effective in investigating mechanisms of proarrhythmia, including those associated with intracellular calcium handling [26, 27, 32, 33]. Variability was considered by modulating the maximal densities of the main transmembrane ionic currents and intracellular fluxes regulating the AP and calcium handling in the mouse ventricular cardiomyocyte discussed as follows. An initial population of 1000 models, representing 1000 cells with different ionic properties, was generated by means of uniform Latin hypercube sampling of scaling factors (scaling from half to two fold) for the following model components: I_CaL_, as main driver of calcium influx into the cardiomyocyte; SR calcium release flux from RyR (J_RyR_), maximal lysosomal calcium uptake and release fluxes (*j*_*clc*_ and *j*_*tpc*_), including calcium leak fluxes (*j*_*clc,leak*_ and *j*_*tpc,leak*_); Na^+^/Ca^2+^ exchanger as principal mediator of calcium extrusion (I_NCX_); SR calcium reuptake through SERCA (J_SERCA_); and the transient outward K^+^ channel (I_to_) and rapid delayed rectifier K^+^ channel (I_Kr_), as main repolarisation currents in mouse ventricular cardiomyocytes. Models were accepted in the WT population if they did not exhibit spontaneous calcium release in cytosolic calcium in any of our baseline protocols (CTRL, NAADP-AM, and ISO), resulting into a total of 343 accepted models, while covering a wide range of cellular profiles and biomarkers. The above acceptance criteria allowed us to investigate the role of lysosomal calcium handling under normal (non-proarrhythmic) physiological function, as well as to dissect its contribution under specific proarrhythmic conditions.

### Quantitative analysis of ionic and calcium features

Integrals of calcium fluxes and concentrations were computed for the entire beat duration using the trapezoidal rule. Amplitude biomarkers were calculated as the absolute difference of maximum and minimum values. Boxplots are used to present the median (central line), 25^th^ and 75^th^ percentiles (box limits) of a distribution. Whiskers extend to the most extreme data points not considered as outliers, and outliers are depicted individually as independent data points. Fold changes of the calcium-related amplitudes were calculated from the medians of the populations being compared. When distributions are shown as line plots, the central line represents the mean, and the error bar limits are the standard error.

## Results

### Model validation of lysosomal calcium handling

To investigate the role of lysosomal calcium handling on ventricular function, the mouse ventricular model of Morotti *et al*. [24] was extended to include a lysosomal compartment (see Methods). In brief, based on experimental evidence on organelle localisation and function [18, 19, 28, 29], lysosome action was modelled as an additional calcium pool, with function primarily driven by calcium loading from the junctional subspace, and NAADP-modulated release to the cytosol via TPC2 channels. The lysosome pool can thereby act as an additional calcium source depending on lysosomal calcium concentration, contributing to the CICR process.

Lysosome model calibration was conducted as follows. Without activating ligands, NAADP-modulated TPC2 openings are typically very brief, with open probability *P*_0_ < 0.0009 [34]. These were described by adopting a previously validated Gaussian distribution on NAADP concentration and an endogenous *[NAADP]* value of 1 nM [30], yielding an upper bound for *P*_0_ of 0.0016, close to the experimental findings. *[NAADP]* was increased to 15 nM when simulating fully opened TPC2 channels, resulting in a *P*_0_ = 0.0134, close to the maximum open probability observed in experiments [34].

The magnitudes of lysosomal calcium fluxes were then calibrated based on experimental data from both wild-type (WT) and genetically-modified TPC2-KO mouse cardiomyocytes [28]. In these experiments (see Figure 2), application of exogenous NAADP (NAADP-AM) increased calcium transient amplitude in WT, but not in TPC2-KO cardiomyocytes. Simulated NAADP-AM action by increased *[NAADP]* resulted in a stronger calcium release from the lysosome compartment via TPC2, recapitulating the increase of cytosolic calcium amplitude (Figure 2A). When lysosomal calcium release (*J*_*ls,rel*_) was set to zero (denoted as TPC2-KO) to mimic the genetically-modified TPC2-KO cardiomyocytes, cytosolic calcium transient features were unaltered by an increase in *[NAADP]*, in agreement with the experimental data (Figure 2B).

Further validation of the lysosomal calcium handling model was conducted under additional pharmacological protocols exerting an action on the CICR process. Experimentally, β-adrenergic stimulation by isoprenaline (ISO) increased calcium transient amplitude in both WT and TPC2-KO, with the response of TPC2-KO significantly reduced compared to WT (Figure 2C-2D). Both cardiomyocyte types also showed faster calcium transient relaxation in response to ISO. Likewise, the activation of the β-adrenergic response in the model by application of 100 nM ISO replicated these effects, with a larger response in WT.

### Lysosomal calcium release primarily modulates intracellular calcium handling by increasing SERCA reuptake and RyR release

In brief, our compartmentalised model of lysosomal calcium handling acts as a fast calcium buffer, enabling an additional direct transport of calcium between the junctional and cytosolic spaces. In the absence of such a buffering effect, junctional calcium can only flow into the cytosol by pure intracellular diffusion, which is characterised by slower transport kinetics.

To investigate how these buffering effects modulate intracellular calcium handling at the whole cellular level, a population of models approach was exploited to account for cellular variability in ionic currents and intracellular calcium fluxes, and their modulation of the full CICR process [26, 27]. To further enable investigations of the role of lysosomal calcium handling on arrhythmogenesis, only models characteristic of healthy physiological function (i.e., not exhibiting spontaneous calcium arrhythmic triggers in any of the described simulated pharmacological protocols) were retained in the population, giving a total of 343 accepted models (see Methods). Subsequent data are reported using this final population.

The boxplots in Figure 3 show the amplitude distributions of calcium concentrations in the lysosome (Figure 3Ai) and cytosolic compartments (Figure 3Bi), and the intracellular calcium fluxes of SERCA reuptake (Figure 3Ci) and RyR release (Figure 3Di) involved in intracellular calcium handling. Representative traces are shown for basal conditions (CTRL) and the investigated protocols of increased lysosomal function (NAADP-AM and ISO), as well as for WT vs TPC2-KO cardiomyocytes (Figures 3Aii-3Dii, 3Aiii-3Diii). For CTRL, lysosomal calcium concentration was higher in TPC2-KO than WT, as a result of the modelled genetic deletion of TPC in TPC2-KO cardiomyocytes (Figures 3Ai-3Aiii), which precludes calcium from being released from the lysosomal compartment. No main median differences in cytosolic calcium amplitude were observed between WT and TPC2-KO (Figures 3Bi-3Biii) in CTRL, consistent with non-significant experimental differences in calcium transient amplitude between both cardiomyocyte types in basal conditions [28]. Evaluation of SERCA and RyR amplitudes showed no main median differences in CTRL (Figures 3Ci-3Ciii, 3Di-3Diii).

**Figure 3.**
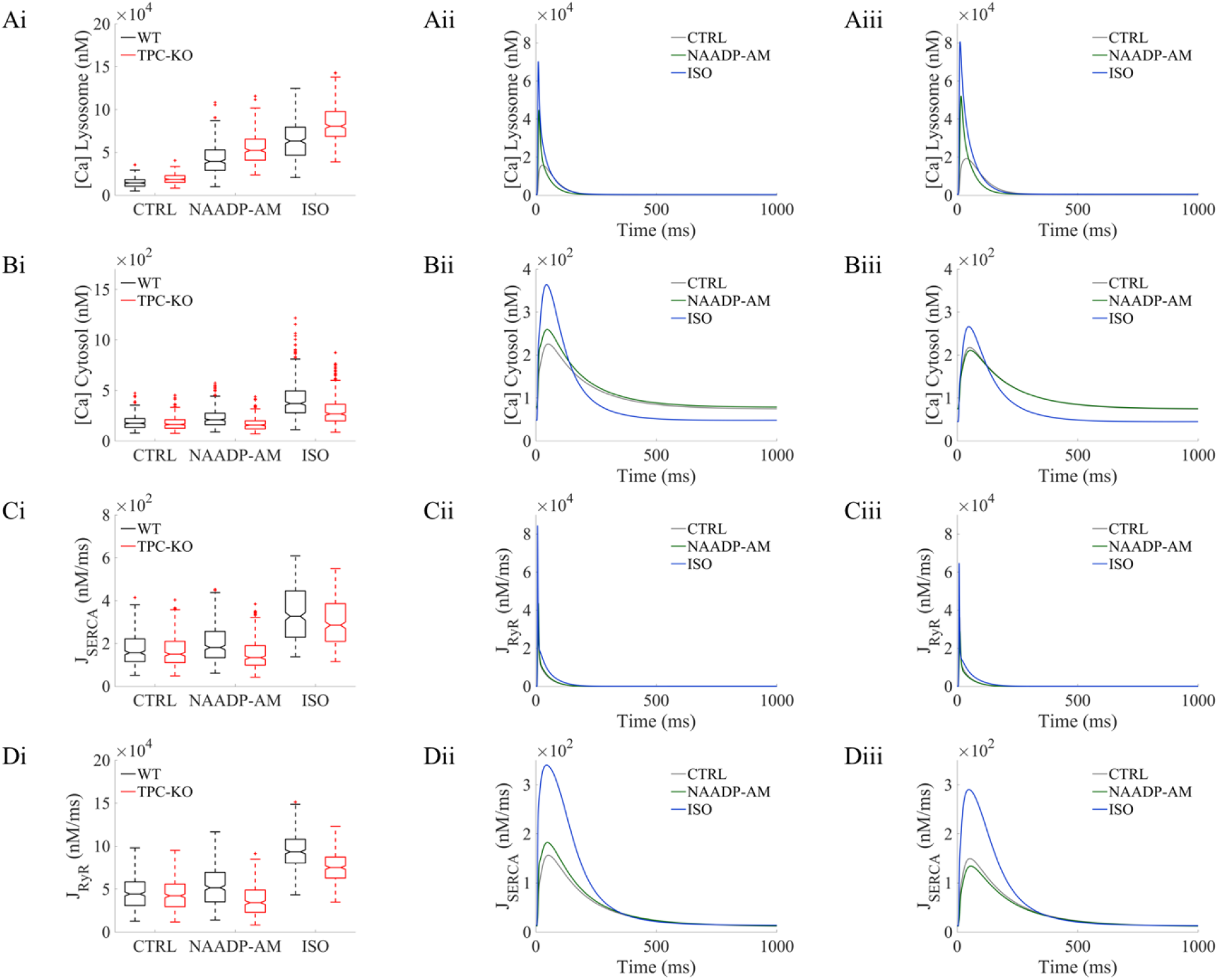
Lysosomal calcium release primarily modulates intracellular calcium handling by increasing SERCA reuptake and RyR release. Model outputs for steady-state are shown for: lysosomal calcium concentration (Ai-iii), cytosolic calcium concentration (Bi-iii), SERCA reuptake (Ci-iii) and RyR release (Di-iii). For each quantity, the distribution of amplitudes under each experimental condition is shown in (i). Wild-type, WT, data is presented in black and TPC2 knock-out, TPC2-KO, in red. The central line represents the median, boxes the inter-quartile range and dotted lines the range. Each quantity is reported under control conditions (CTRL), in the presence of NAADP (NAADP-AM, 15 nM) and in the presence of β-adrenergic stimulation with isoprenaline (ISO, 100 nM). Representative traces of model outputs for WT and TPC2-KO are presented in (ii) and (iii) respectively (CTRL in black, NAAP-AM, in blue, ISO in green).

Under application of exogeneous NAADP (NAADP-AM protocol), the increased lysosomal calcium loading and the increased lysosomal calcium release led to a 1.21-fold amplitude increase of cytosolic calcium in WT (Figure 3Bi). This difference was also apparent in the representative traces of cytosolic calcium concentration (Figure 3Bii). In particular, these effects led to a systolic increase in cytosolic calcium and similar diastolic levels in WT cardiomyocytes under NAADP-AM compared to CTRL (Figure 3Bii), while the cytosolic calcium transient was unaltered upon NAADP-AM application in TPC2-KO cardiomyocytes (Figure 3Biii). Quantitatively, the calibrated population predicted a normalised increase in cytosolic calcium amplitude under NAADP-AM relative to CTRL of 0.208±0.003 (mean ± standard error) in WT vs –0.048±0.000 in TPC2-KO. These values are in agreement with the ranges of 0.16±0.05 for WT and –0.06±0.04 for TPC2-KO in experimental findings [28], further validating model predictions. Mechanistically, the increased cytosolic calcium loading under NAADP-AM in WT cells consequently promoted calcium reuptake by SERCA, causing a 1.16-fold increase in SERCA amplitude (Figure 3Ci), which was also linked to a 1.11-fold increase calcium release by RyR (Figure 3Di). No main median differences with respect to basal conditions were observed in the amplitude of these fluxes for TPC2-KO cardiomyocytes.

When β-adrenergic stimulation by ISO was simulated, the activation of PKA phosphorylation targets (I_CaL_, RyR, phospholamban, and phospholemman in our model) resulted in a higher lysosomal calcium load and release compared to the NAADP-AM protocol (Figure 3Ai). Matching the experimental findings, there was also a greater response to ISO in WT cells than in TPC2-KO; isoprenaline application increased the cytosolic calcium amplitude by 2.14-fold compared to CTRL for WT, and by 1.65-fold in WT versus TPC2-KO models (Figure 3Bi). In terms of normalised changes of cytosolic calcium amplitude, the population predicted increases of 1.183±0.028 in WT and 0.675±0.020 in TPC2-KO, are in agreement with the ranges of 1.23±0.24 and 0.57±0.17 increase in experimental findings, respectively [28]. Isoprenaline application led to a stronger 2.09-fold increase in calcium reuptake by SERCA in WT versus TPC2-KO (Figure 3Ci), and a 2.11-fold increase of calcium release by RyR (Figure 3Di).

Altogether, the results presented in this section qualitatively and quantitatively recapitulate the findings associated with experimental observations of lysosomal function on the calcium transient in both WT and genetically-modified TPC2-KO mouse cardiomyocytes [28]. These results also illustrate an impact of lysosomal calcium in intracellular calcium handling by increasing the cytosolic calcium peak concentration, which in turn potentiates SERCA reuptake, increases the SR calcium load, and promotes CICR via RyR release.

### Lysosomal calcium release promotes spontaneous calcium release under hypercalcemia and β-adrenergic stimulation by increasing junctional and SR calcium gradient

Following our model validation and mechanistic findings of a lysosomal modulation of CICR under normal calcium loading, we further tested the model to investigate mechanisms of lysosomal involvement under proarrhythmic conditions.

Firstly, we simulated conditions of calcium overload by hypercalcemia (raising extracellular calcium from 1.0 to 1.8 mM), coupled with β-adrenergic stimulation. Out of the 343 calibrated cellular profiles (none exhibiting spontaneous release events in basal conditions), 57% (194/343) of the models displayed spontaneous calcium release events in WT compared to 45% (155/343) in TPC2-KO models, highlighting a lysosomal involvement in the triggering of intracellular calcium proarrhythmic events.

To better understand the mechanisms where lysosomal calcium handling is directly responsible for these proarrhythmic events, TPC-specific proarrhythmic profiles (n=39) were extracted from the population, meaning that the selected models exhibited spontaneous calcium release in WT (i.e., with active lysosomal TPC release) but not in their paired TPC2-KO configurations. In terms of ionic properties underlying cell-to-cell variability, proarrhythmic profiles featured a trend of favouring larger lysosomal calcium uptake than release, with the largest difference exhibited by TPC2-specific proarrhythmic models (J_ls,up_ and J_ls,rel_, Figure S1A). Proarrhythmic models also displayed a trend towards a larger SR calcium release than uptake (J_SERCA_ and J_RyR_, Figure S1A), which was reversed in the non-proarrhythmic profiles. Importantly, the TPC-specific proarrhythmic profiles were characterised by the smallest densities in L-type calcium current, while this was the largest in the rest of proarrhythmic profiles (I_CaL_, Figure S1A), as also verified at the level of total (integral) current (Figure S1B). Taken together, our findings highlight that, in the TPC-specific proarrhythmic models, spontaneous calcium release events were directly mediated by lysosomal calcium concentration: their intermediate lysosomal calcium load (sitting between that of other proarrhythmic and non-proarrhythmic models; Figure S1C) provides the sufficient additional calcium load to spark the event, compensating their smaller SR concentration and calcium release via RyR (Figures S1E, S1F). On the other hand, spontaneous calcium release events in non-TPC-specific proarrhythmic profiles were primarily due to the larger I_CaL_ and resulting SR calcium accumulation (Figures S1B, S1E), irrespective of any lysosomal contribution, as the spontaneous events also occurred when blocking lysosomal release (*J*_*ls,rel*_ = 0). Finally, non-proarrhythmic profiles compensated their intermediate I_CaL_ (Figure S1B) through their trend towards larger SR calcium uptake than release (Figures S1A, S1D, S1E), minimising the difference in calcium concentration as driver for spontaneous release between the junctional and SR subspaces (Figure S1H).

A further analysis of lysosome modulation of spontaneous calcium release events is presented in Figures 4 and 5. Figure 4 illustrates the traces of the main calcium concentrations and fluxes involved in lysosomal calcium release under hypercalcemia and β-adrenergic stimulation for the TPC-specific proarrhythmic profiles. Figure 5 quantitatively summarises the relevant metrics as integrals over an entire beat. Hypercalcemia resulted in lysosomal overload (Figure 4Ai) compared to basal conditions (Figure 3Ai), which contributed to a stronger calcium release from lysosome to cytosol (Figure 4Bi). This was confirmed by an increase of total calcium reuptake by SERCA (Figures 4Ci, 5A), leading to a higher SR calcium concentration and consequently increasing the likelihood of spontaneous events at RyR (Figures 4Di, 5B).

**Figure 4.**
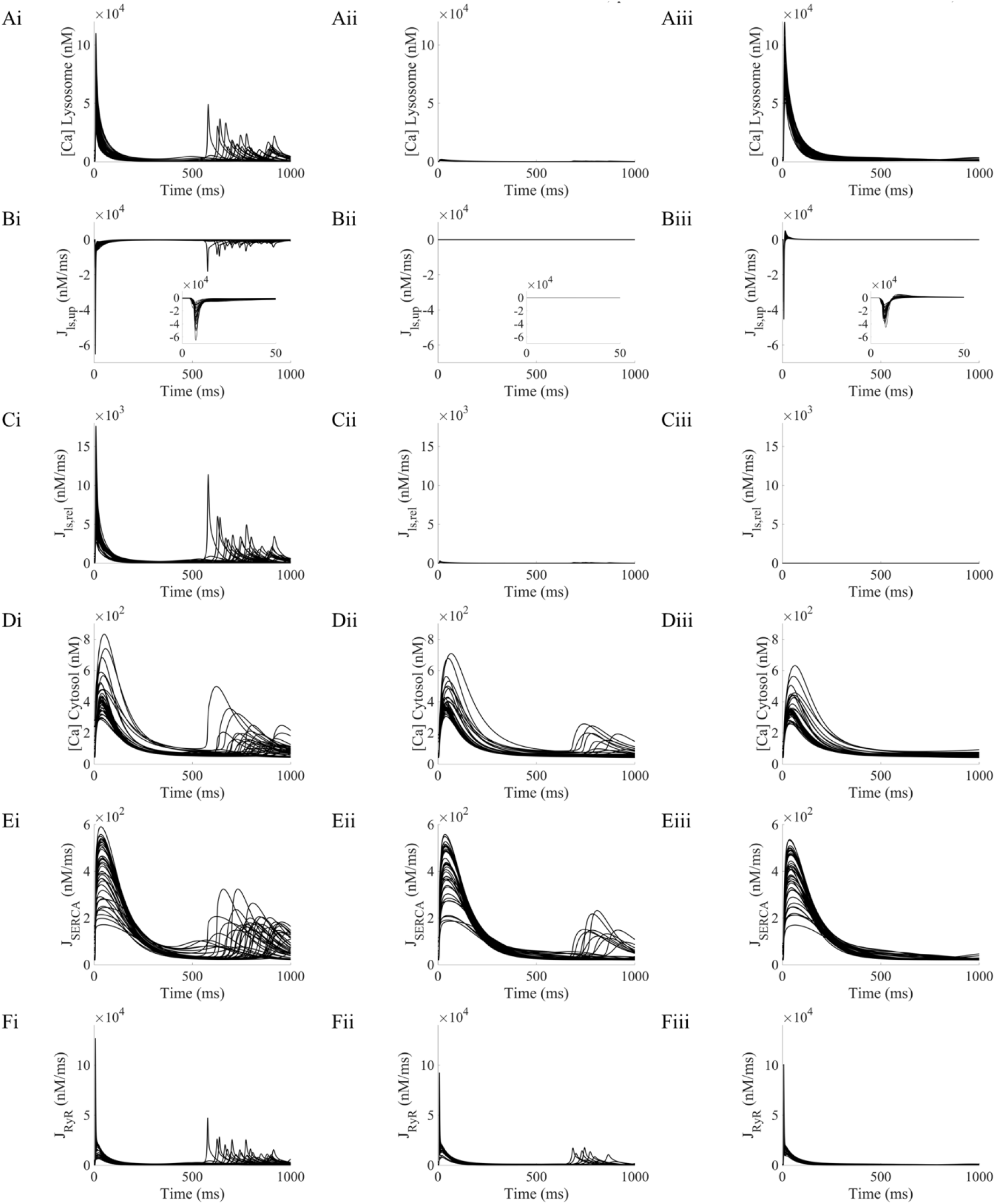
Lysosomal calcium release promotes spontaneous calcium release in TPC-specific pro-arrhythmic profiles under hypercalcemia and β-adrenergic stimulation by increasing the junctional-SR calcium gradient. **4Ai-4Fi:** Lysosomal calcium concentration, lysosomal uptake and release, cytosolic calcium concentration, and SR reuptake and release fluxes, respectively, in basal conditions. **4Aii-4Fii:** Calcium concentrations and fluxes under lysosomal uptake block (*J*_*ls,up*_ = 0). **4Aiii-4Fiii:** Calcium concentrations and fluxes under lysosomal release block (*J*_*ls,rel*_ = 0).

**Figure 5.**
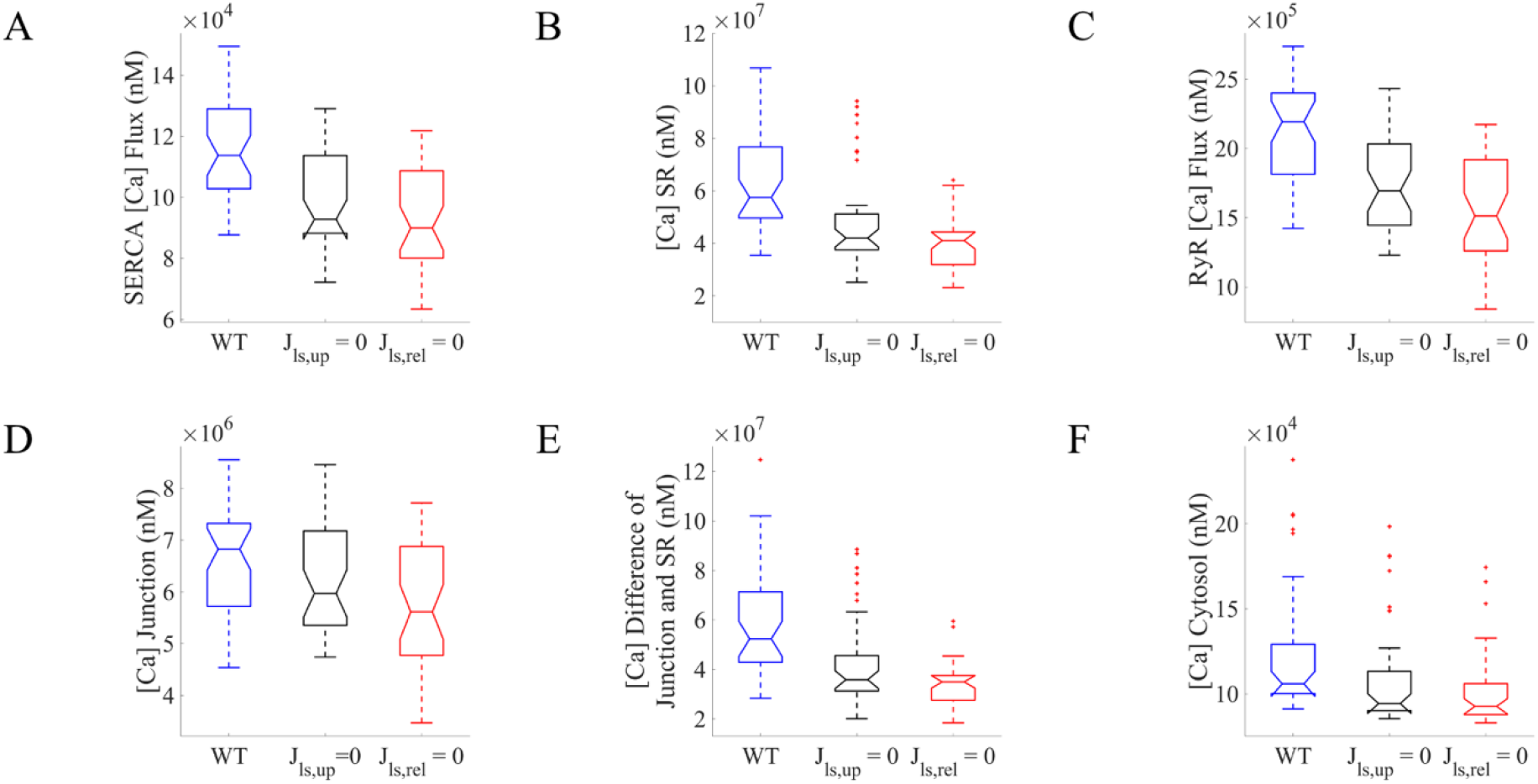
Loss of each lysosomal flux reduced spontaneous calcium release from SR in TPC-specific pro-arrhythmic models under hypercalcemia and β-adrenergic stimulation. **5A-5F:** Distributions of total (integral) calcium fluxes and concentrations over a complete beat, comparing scenarios in WT (blue), blocking lysosomal calcium loading (*J*_*ls,up*_ = 0, black), and blocking lysosomal calcium release (*J*_*ls,rel*_ = 0, red).

Continuing to focus on models with TPC-dependent arrhythmias, the influence of lysosomal calcium handling in mediating spontaneous calcium release events was further dissected by blocking each of the respective lysosomal fluxes in the WT. The block of lysosomal calcium uptake (*J*_*ls,up*_) precluded calcium loading in the lysosome compartment (Figure 4Aii), reducing spontaneous calcium release events to 26% (10/39) compared to TPC-specific pro-arrhythmic models in WT (Figure 4Dii). The models no longer exhibiting spontaneous calcium release events were characterised by smaller lysosomal uptake and release fluxes, in spite of exhibiting a stronger SR reuptake and release, but a weaker magnitude of the L-type calcium current (Figures S2A-S2B). In terms of impact on function, the block of the lysosomal uptake flux impeded the loading of the lysosomal compartment, resulting in the loss of lysosomal calcium release (Figure 4Aii-4Cii). The absence of the fast buffering calcium effect via the lysosomes decreased calcium transient amplitude (Figure 4Dii) and thus calcium reuptake by SERCA (Figures 4Eii, 5A), decreasing SR concentration (Figure 5B), and therefore the magnitude of RyR release (Figures 4Fii, 5C). This in turn decreases the junctional calcium concentration (Figure 5D) and the gradient of calcium concentrations between the junction and SR (Figure 5E), as the main driving force for spontaneous calcium release once RyR recovers excitability.

On the other hand, loss of the lysosomal release in TPC2-KO cardiomyocytes (as mimicked by setting *J*_*ls,rel*_ = 0) had a dual anti-arrhythmic effect, resulting in the complete abolishment of spontaneous calcium release events in all TPC-specific pro-arrhythmic models. First, although the lysosome compartment now gets loaded (Figure 4Aiii), the block of lysosomal release precludes its calcium load from flowing directly into the cytosol, getting transiently stored for a longer period of time in this compartment. The absence of the fast buffering lysosomal pathway thus resulted in a similar anti-arrhythmic mechanism as above described in the case of lysosomal uptake block (decreased calcium transient amplitude, Figure 4Diii; reduced SERCA uptake, Figures 4Eiii, 5A; reduced SR concentration, Figure 5B; and therefore, a smaller magnitude of RyR release, Figures 4Fiii, 5C). Second and even more importantly, part of the lysosomal calcium concentration (approximately 83% in median) is reverted into the junctional space through the lysosomal loading channels, which is even noticeable in steady-state conditions (positive peak in Figure 4Biii). The additional calcium being released back into the junction contributes to a reduced driving force for CICR, which is translated in the long term to reduced calcium loads in the cytosol, SR, and junctional spaces (Figures 5B, 5D, 5F), drastically diminishing the driving force for ion flow (Figure 5E), and therefore for spontaneous calcium release, when RyRs open.

In summary, approximately 20% (39/194) of our investigated models exhibiting spontaneous calcium release events under hypercalcemia and β-adrenergic stimulation were directly linked to lysosomal action, due to an increased calcium gradient between the junctional space and SR. Such lysosome-specific events can be effectively counteracted by inhibition of lysosomal calcium release, in agreement with experimental findings of decreased arrhythmic propensity in TPC2-KO mice [28].

### Lysosomal calcium release promotes spontaneous calcium release under fast pacing and β-adrenergic stimulation by increasing junctional and SR calcium gradient

Following experimental evidence of an involvement of lysosomal calcium handling in mediating arrhythmic events under chronic β-adrenergic stimulation and burst pacing [28], we hypothesised similar proarrhythmic mechanisms due to intracellular calcium accumulation by fast pacing as in concomitant hypercalcemia and sustained β-adrenergic stimulation. For these investigations, our populations of cardiomyocyte models were paced at 10 Hz for 150 beats to raise intracellular calcium, followed by a transition to slow pacing at 1 Hz for 5 consecutive beats, where spontaneous calcium release events were recorded.

Out of the 343 calibrated cellular profiles, 63% (217/343) of WT models displayed spontaneous calcium release events during their transitions to slow pacing, compared to 46% (159/343) in TPC2-KO models, evidencing a lysosomal contribution to intracellular calcium proarrhythmic events. TPC-specific proarrhythmic profiles (n=58) were extracted from the new population, based on those exhibiting spontaneous calcium release events when transitioning to slow pacing in WT, while not in their paired TPC2-KO configuration. In terms of ionic properties underlying cellular variability, TPC-specific proarrhythmic profiles were characterised by larger I_CaL_ and I_NCX_ densities, together with intermediate levels of SERCA reuptake (J_SERCA_) and RyR release (J_RyR_) (Figure S3A). The rest of the proarrhythmic models followed a clear trend towards stronger SR release than uptake, which was only reverted in the non-proarrhythmic profiles (Figure S3A).

The key intracellular concentrations and calcium fluxes involved in CICR are presented in Figure 6 for the TPC-specific proarrhythmic models during their transition to slow pacing. In WT, intracellular calcium accumulation due to the fast pacing portion of the protocol is apparent at the start of the transition (Figure 6Di), compared to basal conditions (Figure 3Bi). This calcium overload is taken back up into the SR by SERCA (Figure 6Ei), leading to subsequent SR overload and a persistent release by RyR in the form of spontaneous calcium events during the entire transition in pacing (Figure 6Fi). Fast pacing model runs were repeated at a faster pacing rate of 25 Hz and analogous findings were observed (Figure S4).

**Figure 6.**
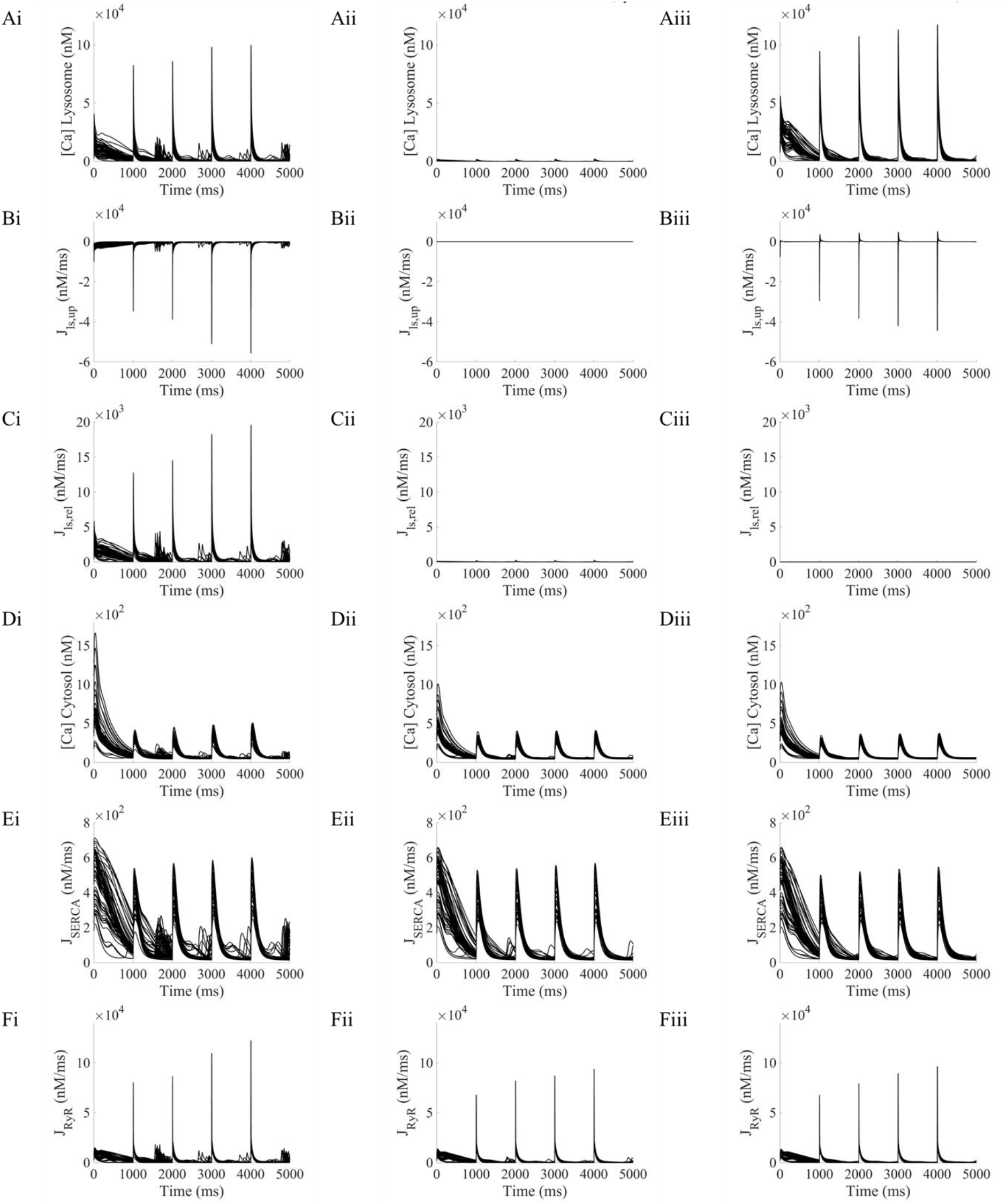
Lysosomal calcium release promotes spontaneous calcium release in TPC-specific pro-arrhythmic profiles under fast pacing and β-adrenergic stimulation, by increasing the junctional-SR calcium gradient. All results presented under initial fast pacing at 10 Hz. Cessation of pacing occurred at time 0 (referred as by beat 0). Analyses contributing to further figures are taken from the first four beats at 1 Hz, ie. 1000-5000 ms. **6Ai-6Fi:** Lysosomal calcium concentration, lysosomal uptake and release, cytosolic calcium concentration, and SR reuptake and release fluxes, respectively, in basal conditions. **6Aii-6Fii:** Calcium concentrations and fluxes under lysosomal uptake block (*J*_*ls,up*_ = 0). **6Aiii-6Fiii:** Calcium concentrations and fluxes under lysosomal release block (*J*_*ls,rel*_ = 0).

The influence of lysosomal calcium handling in mediating these spontaneous calcium release events was investigated by blocking each of the respective lysosomal fluxes. Figure 7 shows distributions of total (integral) calcium fluxes after cessation of fast pacing. While for the hypercalcemia case simulations were in a stable state, the cessation of fast pacing produced beat-by-beat changes, so results are presented as mean±SD to assist visualization and avoid multiple unconnected boxplots. Similar to the hypercalcemia case, block of the lysosomal uptake (*J*_*ls,up*_ = 0) precluded calcium load in the lysosome (Figures 6Aii-6Cii), leading to the abolition of the fast buffering calcium pathway from the junctional space into the cytosol. This resulted into the previously discussed cascade of decreased cytosolic calcium transient amplitude (Figures 6Dii, 7F), reduced SERCA uptake (Figures 6Eii, 7A), decreased SR loading (Figure 7B), and smaller magnitude of the junction-SR calcium gradient as driving force for RyR release (Figures 6Fii, 7I, 7K), decreasing the incidence of spontaneous calcium release events to only 21% (12/58) of the TPC-specific proarrhythmic models. Block of the lysosomal release (*J*_*ls,rel*_ = 0) was accompanied by an additional reversal of part of the lysosomal calcium concentration into the junctional space via the lysosomal loading channel (positive peaks in Figure 6Biii), further decreasing the junction-SR gradient for spontaneous RyR release (Figure 7E). Although this reduction was markedly smaller than in the hypercalcemia case, it yet sufficed to completely abolish all spontaneous calcium release events in the TPC2-specific proarrhythmic models. The same findings were obtained under faster pacing rates (Figure S5).

**Figure 7.**
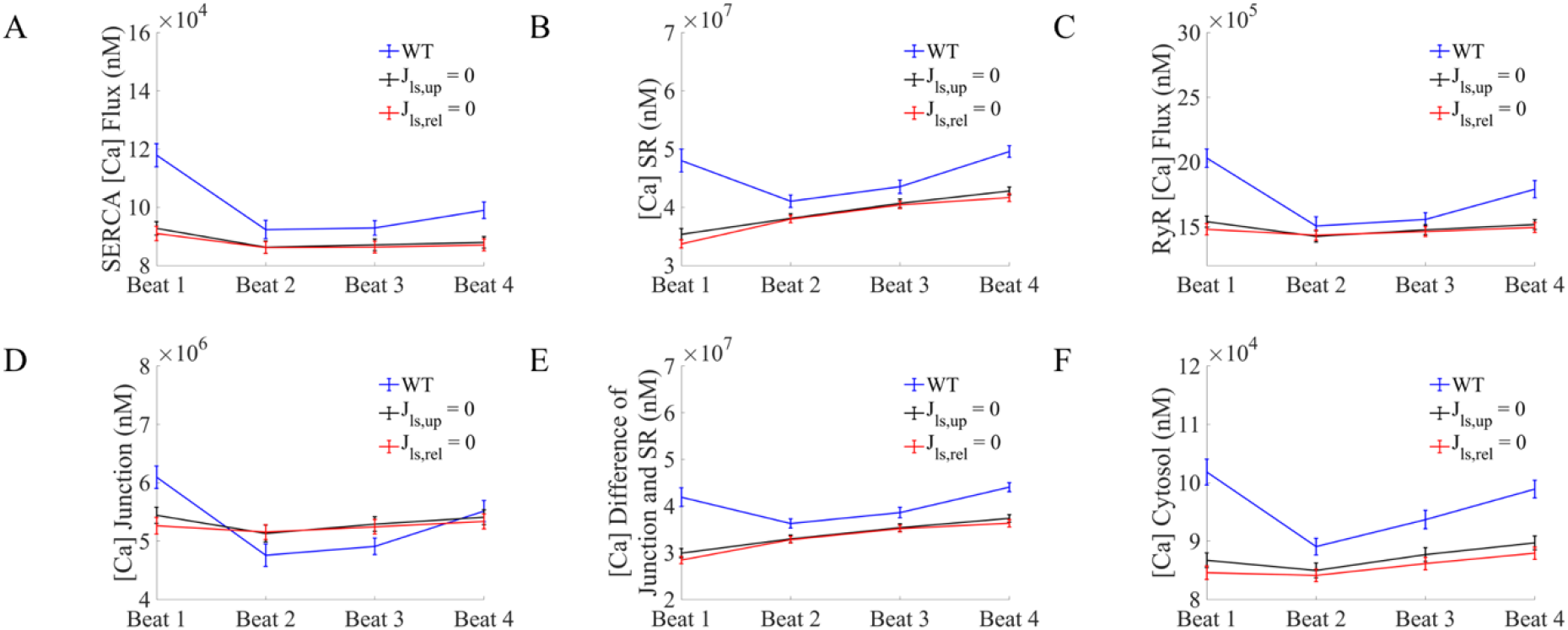
Loss of each lysosomal flux reduced spontaneous calcium release from SR in TPC-specific pro-arrhythmic models under fast pacing and β-adrenergic stimulation. **7A-7F:** Distributions of total (integral) calcium fluxes and concentrations per beat in the transition to slow pacing (data presented as mean±SD), comparing scenarios in WT, blocking lysosomal calcium loading (*J*_*ls,up*_ = 0), and blocking lysosomal calcium release (*J*_*ls,rel*_ = 0). Beats are numbered from beat 0 (not shown, as state is representative of fast pacing). All results presented under initial fast pacing at 10 Hz.

## Discussion

In this study we present modifications to the well-established Morotti *et al*. cardiomyocyte model [24] to include a lysosomal calcium pool. To our knowledge, this work represents the first cardiac computational model to include a lysosomal compartment as part of beat-to-beat calcium signalling. Lysosomes, historically thought of as waste processing units, are now accepted to be complex signalling organelles. They are a store of calcium [35] and release calcium in response to the endogenous second-messenger NAADP [16]. Work published from our group and others has confirmed that lysosomal calcium signalling is constitutively active in both ventricular [20, 21] and atrial [12] cardiomyocytes and contributes to β-adrenergic responses [12, 20, 21, 28, 36]. Further, abolishing lysosomal calcium signalling, either pharmacologically or by genetic ablation of the NAADP effector channel TPC2, is protective against cardiac hypertrophy and arrhythmias [21, 28]. No cardiomyocyte model to date has sought to include lysosomal calcium signalling and we therefore set out to add a lysosomal calcium pool to a previously published, widely accepted model of mouse ventricular electrophysiology. This modified model was used to probe the impacts of lysosomal calcium signalling under both physiological and pro-arrhythmic conditions. After initial model construction, testing and validation (Figure 2), we used a population of models approach to generate a pool of models with a range of expression levels in other signalling proteins (e.g., I_CaL_, RyR, SERCA). Our model population was able to recapitulate experimental data: ‘wild-type’ cells responded to NAADP with an increase in calcium transient amplitude, TPC2KO cells did not (Figure 3); TPC2KO cells exhibited a reduced response to β-adrenergic stimulation by ISO (Figure 3) and a reduced propensity to arrhythmogenesis under pro-arrhythmic conditions (Figures 4 and 6). The major results of our simulation studies are consistent with a role for lysosomes in increasing calcium uptake to the SR by SERCA, release from the SR by RyRs and SR-junction concentration gradient (Figures S1 and S3). The overall effect is to increase calcium release from the SR during calcium-induced calcium release. In a small fraction of the models in our population, lysosomal calcium was found to be the direct cause of arrhythmogenesis under pro-arrhythmic conditions. Further consideration of these models showed that they did not exhibit spontaneous lysosomal release but delayed after-depolarisations arising as a result of RyR-mediated spontaneous calcium release from the SR. Our further trials (Figures 5, 7 and S2) suggested that, when the balance of other signalling effectors is appropriate, lysosomes acting as a fast buffer can effectively shunt the location of calcium to promote delayed afterdepolarisation formation.

The Morotti *et al*. model [24] was chosen for modification to include the lysosomal calcium pool because it is closest in nature to our published laboratory research [28]; it is a murine ventricular cell model detailing both calcium and action potential characteristics. The Morotti *et al*. model also includes validated β-adrenergic and CaMKII signalling modules, both of which have been implicated in NAADP-mediated signalling in the heart [28, 37]. The updated model presented in this paper was validated using published murine single cell and ex-vivo tissue data from our group [28] with the position of the lysosomal calcium pool justified by our structural studies [19]. The lysosomal pool features calcium uptake, calcium release and calcium leak. The calcium release mechanism is via TPC2 channels, with the TPC2 open probability in response to NAADP modelled on the basis of data from Penny *et al*. [30]. 15 nM NAADP-AM was used throughout our study, as the NAADP concentration equivalent to nearly maximal TPC2 open probability.

The addition of the lysosomal calcium pool to our model also included a calcium leak channel and a calcium uptake mechanism. Although the candidate protein for neither of these processes has been identified, it is justified to include them here: calcium leak must occur from lysosomes as breakdown of the lysosomal hydrogen ion gradient following treatment with Bafilomycin-A1 leads to a run-down of lysosomal calcium in the absence of other stimulation [35, 38]. Similarly, there must be a calcium uptake mechanism, as the lumenal calcium of lysosomes is several orders of magnitude higher than the cytosolic concentration [35]. In the absence of characterised mechanisms for this process we adopted an analogous formulation to TPC2 channels. This correctly recapitulated our calcium transient data under different pharmacological protocols. This modelling assumption can nevertheless be easily revisited, shall additional datasets on lysosomal calcium loading become available.

The study presented in this paper used a population-of-models approach to take into account that cells are not a homogenous population and may respond to differing degrees based on inherent expression/activity level of various known calcium signalling and electrophysiology components. To this end, Latin hypercube sampling was used to create a population of 1000 models to represent a wide combination of underlying contributions. Of these, a sub-population of 343 were accepted into the final population as behaving within the physiological envelope of our experimental data; they were not arrhythmic under any ‘physiological’ conditions. In our simulations using this population of models, lysosomal calcium release primarily modulated intracellular calcium handling by increasing SERCA reuptake and subsequently RyR release. This is consistent with published data which suggest that stimulation of lysosomal calcium release with NAADP leads to an increase in SR calcium load, as measured by caffeine transient [12, 20]. The underlying action of the lysosomal calcium pool was essentially to act as a fast buffer, moving calcium into the space from which SERCA pumps calcium into the SR. In comparison to WT, TPC2-KO models exhibited reduced cytosolic calcium concentration, SR calcium uptake (J_SERCA_) and RyR calcium release (J_RyR_) upon application of NAADP or ISO.

Our published work in ventricular cardiomyocytes and ex vivo cardiac tissue suggest that TPC2-KO is protective against the development of cardiac arrhythmias under conditions of acute or chronic high-dose β-adrenergic stimulation [28]. This is also true of acute ISO exposure in single cells where TPC2 channels have been inhibited pharmacologically [21]. As an analogous set of experiments, we exposed the population of 343 accepted models to pro-arrhythmic conditions in both WT and TPC2-KO configurations. Synergetic to β-adrenergic stimulation, these conditions were either raised extracellular calcium or rapid burst pacing, both of which are common proarrhythmic interventions in experimental studies of calcium overload. Similar to published data, we found that TPC2-KO is protective against the genesis of arrhythmia: 57% of WT models exhibited spontaneous events under β-adrenergic stimulation and high calcium, but this fell to 45% in TPC2-KO. Moreover, 63% of WT models exhibited spontaneous events under β-adrenergic stimulation and fast pacing, but this fell to 46% in TPC2-KO.

In a sub-population of the accepted models exposed to pro-arrhythmic conditions, the WT model was found to be arrhythmic and the corresponding TPC2-KO was not. That is to say the only difference between the rhythmic and arrhythmic simulations was the presence of lysosomal calcium signalling. These models comprised n=39 for raised extracellular calcium with β-adrenergic stimulation and n=58 for rapid pacing and β-adrenergic stimulation, or 20% (39/194) and 27% (58/217) of all models which exhibited arrhythmia for each pro-arrhythmic condition respectively. We looked further at these TPC2-dependent arrhythmia models to try to determine what the action(s) of the lysosome might be under these circumstances, and therefore by which specific mechanism lysosomal calcium might be contributing to the genesis of rhythm disturbance under our pro-arrhythmic conditions, and/or in what way TPC2-KO is protective against arrhythmia. Analysis of the intracellular calcium traces during the genesis of spontaneous activity showed a profile of DADs arising from spontaneous SR calcium release (Figures 4 and 6). In other cell types, from various species, lysosomal calcium release can lead to a cellular calcium transient, or cellular calcium oscillations, directly by recruiting calcium-induced calcium release from the reticular store [16, 39, 40]. There were, however, no instances in which we observed something akin to spontaneous lysosomal calcium release leading directly to a cytosolic calcium transient. This is consistent with published experimental work, which demonstrates the role of lysosomal calcium signalling in cardiomyocytes as increasing the size of the normal, electrically-stimulated calcium transient as opposed to generating de-novo transients by CICR [12, 20, 28]. Although Nebel and co-authors reported the generation of a spontaneous signal on patch-application of NAADP [21], we have not observed any similar phenomena through application from the patch pipette (unpublished research), ultraviolet photolysis of a caged compound [12, 20, 28] or via the membrane using an esterified form [12, 28]. The results seen in our simulation study therefore also support our published experimental findings.

One way to consider the specific effects of lysosomal fluxes on models exhibiting TPC2-dependent arrhythmias is to investigate the consequence of removing lysosomal calcium contributions, flux-by-flux, from the WT versions of the models in question. Removing these fluxes from the models with TPC2-dependent arrhythmias (Figures 5 and 7), regardless of the specific pro-arrhythmic condition, led to decreases in: SERCA calcium flux, RyR calcium flux, SR calcium content and the concentration gradient between the SR and junction. Taken together, this would suggest a conclusion consistent with that seen in our physiological signalling study: the presence of lysosomal calcium release boosts SR calcium content and release by acting as a fast buffer. Further, under particular pro-arrhythmic conditions, the shifting of calcium can occur to the level where the cell becomes at risk of DADs. This mechanism would also be consistent with published cellular recordings which showed a significant increase in SR calcium content, as measured by caffeine transient, after exposure to NAADP [12, 20]. Of note in this regard, the levels of SR-related calcium signalling components in TPC2-dependent arrhythmic models generally fell in between that for models which did not show arrhythmias and models which showed arrhythmias regardless of the presence of TPC2 (Figures S1 and S3). This supports the conclusion that lysosomal calcium in TPC2-dependent arrhythmia models is acting to push those otherwise physiological cells into the arrhythmic category via actions related to SR calcium signalling.

In conclusion, we have modified an existing cellular model of cardiomyocyte electrophysiology and calcium handling to include a novel calcium pool, the lysosome. This is the first cardiac cellular model to include lysosomal calcium in beat-to-beat regulation. The lysosomal calcium pool, which was unknown at the turn of the century, is now understood to contribute to beat-to-beat calcium handling, response to β-adrenergic stimulation and potentially to arrhythmogenesis. Our simulations support a role for the lysosome as a fast buffer, acting ultimately to shunt calcium into the SR. The population of models approach allowed us to investigate a range of conditions and support that removing lysosomal calcium signalling by TPC2-KO reduces the likelihood of developing spontaneous activity under pro-arrhythmic conditions. Taken another way, our model suggests that there are some conditions under which lysosomal calcium signalling can be pro-arrhythmic. This modelling work leads to an important next experimental step, to investigate what these pro-arrhythmic conditions are in both healthy cells and those in disease states associated with an increased likelihood of arrhythmia.

### Limitations

Our approach for lysosomal calcium handling modelling effectively considers the lysosomes as a separate domain, feeding from the junctional space (shared by I_CaL_ channels and RyRs) and releasing their calcium load into the cytosol. This, together with appropriate considerations of lysosomal volume size, allows the lysosomal compartment to achieve high local calcium concentrations compared to the cytosol. Given that the cytosolic space is shared by CaMKII and phospholamban/SERCA, our lysosomal compartment can therefore be conceptually interpreted as in close proximity to these proteins, with lysosomal calcium release via TPC2 being able to directly modulate their function. Shall sufficient evidence becomes available, the proposed modelling could be expanded to consider different (nano-)domain configurations.

## Author contributions

Zhaozheng Meng: Conceptualization, Methodology, Software, Investigation, Formal analysis, Writing – original draft, review & editing. Rebecca A Capel: Contribution to theoretical development, Writing-review & editing. Samuel J Bose: Results discussion, Writing-review & editing. Erik Bosch: Theoretical discussions, early model development. Sophia de Jong: Theoretical discussions, early model development. Antony Galione: Mechanistic discussions, Writing – review & editing. Robert Planque: Theoretical discussions, early model development, Writing – review & editing. Rebecca-Ann B Burton: Conceptualization, Design & Methodology, Writing – review & editing, Resources, Supervision. Alfonso Bueno-Orovio: Conceptualization, Design & Methodology, Investigation, Writing – review & editing, Resources, Supervision.

## Acknowledgments

This work is supported by Sir Henry Dale Wellcome Trust and Royal Society Fellowship (109371/Z/15/Z; to R.A.B.B.) and an Oxford RCF Award. This project was supported by a British Heart Foundation (BHF) Project Grant (PG/18/4/33521). R.A.C. is a Post-doctoral Scientist funded by the Wellcome Trust and Royal Society (109371/Z/15/Z). S.J.B. is a Post-doctoral Scientist funded by the BHF (PG/18/4/33521). A.B.O. acknowledges a BHF Intermediate Basic Science Fellowship (FS/17/22/32644), and an Impact for Infrastructure Award from the National Centre for the Replacement, Refinement and Reduction of Animals in Research (NC/P001076/1). The authors acknowledge additional support from the Oxford BHF Centre of Research Excellence (RE/13/1/30181), and the use of the University of Oxford Advanced Research Computing (ARC) facility (https://doi.org/10.5281/zenodo.22558). For the purpose of Open Access, the authors have applied a CC BY public copyright licence to any Author Accepted Manuscript (AAM) version arising from this submission. We thank Prof Derek Terrar for sharing his scientific opinions.

## Data availability

All codes underpinning this research are openly available at: https://github.com/zhaomeng132/Ls_mv

## References

1. Nattel, S., B. Burstein, and D. Dobrev, Atrial remodeling and atrial fibrillation: mechanisms and implications. Circ Arrhythm Electrophysiol, 2008. 1(1): p. 62–73.

2. Scoote, M. and A.J. Williams, Myocardial calcium signalling and arrhythmia pathogenesis. Biochem Biophys Res Commun, 2004. 322(4): p. 1286–309.

3. Burton, R.-A.B. and D.A. Terrar, Emerging Evidence for cAMP-calcium Cross Talk in Heart Atrial Nanodomains Where IP_3_-Evoked Calcium Release Stimulates Adenylyl Cyclases. Contact, 2021. 4: p. 1–13.

4. Smith, G.L. and D.A. Eisner, Calcium Buffering in the Heart in Health and Disease. Circulation, 2019. 139(20): p. 2358–2371.

5. Bers, D.M., Cardiac excitation-contraction coupling. Nature, 2002. 415(6868): p. 198–205.

6. Raffaello, A., et al., Calcium at the Center of Cell Signaling: Interplay between Endoplasmic Reticulum, Mitochondria, and Lysosomes. Trends in Biochemical Sciences, 2016. 41(12): p. 1035–1049.

7. Nijholt, K.T., R.A. de Boer, and B.D. Westenbrink, What You Did Not Know About Cardiac Ca2+ Handling Lysosomes and Oxidized PKA. Circulation, 2021. 143(5): p. 466–469.

8. Berridge, M.J., M.D. Bootman, and H.L. Roderick, Calcium signalling: dynamics, homeostasis and remodelling. Nat Rev Mol Cell Biol, 2003. 4(7): p. 517–29.

9. Simon, J.N., et al., Oxidation of Protein Kinase A Regulatory Subunit PKARIα Protects Against Myocardial Ischemia-Reperfusion Injury by Inhibiting Lysosomal-Triggered Calcium Release. Circulation, 2021. 143(5): p. 449–465.

10. Wheat, M.W., Ultrastructure autoradiography and lysosome studies in myocardium. J Mt Sinai Hosp N Y, 1965. 32: p. 107–21.

11. Kottmeier, C.A. and M.W. Wheat, Myocardial lysosomes in experimental atrial septal defects. Circ Res, 1967. 21(1): p. 17–24.

12. Collins, T.P., et al., NAADP influences excitation-contraction coupling by releasing calcium from lysosomes in atrial myocytes. Cell Calcium, 2011. 50(5): p. 449–58.

13. Capel, R.A., et al., Two-pore Channels (TPC2s) and Nicotinic Acid Adenine Dinucleotide Phosphate (NAADP) at Lysosomal-Sarcoplasmic Reticular Junctions Contribute to Acute and Chronic beta-Adrenoceptor Signaling in the Heart. J Biol Chem, 2015. 290(50): p. 30087–98.

14. Wu, P.L., et al., Myocardial Upregulation of Cathepsin D by Ischemic Heart Disease Promotes Autophagic Flux and Protects Against Cardiac Remodeling and Heart Failure. Circulation-Heart Failure, 2017. 10(7).

15. Churchill, G.C. and A. Galione, NAADP induces Ca2+ oscillations via a two-pool mechanism by priming IP3- and cADPR-sensitive Ca2+ stores. EMBO J, 2001. 20(11): p. 2666–71.

16. Churchill, G.C., et al., NAADP mobilizes Ca(2+) from reserve granules, lysosome-related organelles, in sea urchin eggs. Cell, 2002. 111(5): p. 703–8.

17. Galione, A., et al., NAADP as an intracellular messenger regulating lysosomal calcium-release channels. Biochem Soc Trans, 2010. 38(6): p. 1424–31.

18. Ruas, M., et al., Expression of Ca2+-permeable two-pore channels rescues NAADP signalling in TPC-deficient cells. Embo Journal, 2015. 34(13): p. 1743–1758.

19. Aston, D., et al., High resolution structural evidence suggests the Sarcoplasmic Reticulum forms microdomains with Acidic Stores (lysosomes) in the heart. Sci Rep, 2017. 7: p. 40620.

20. Macgregor, A., et al., NAADP controls cross-talk between distinct Ca2+ stores in the heart. J Biol Chem, 2007. 282(20): p. 15302–11.

21. Nebel, M., et al., Nicotinic acid adenine dinucleotide phosphate (NAADP)-mediated calcium signaling and arrhythmias in the heart evoked by β-adrenergic stimulation. J Biol Chem, 2013. 288(22): p. 16017–30.

22. Davidson, S.M., et al., Inhibition of NAADP signalling on reperfusion protects the heart by preventing lethal calcium oscillations via two-pore channel 1 and opening of the mitochondrial permeability transition pore. Cardiovascular Research, 2015. 108(3): p. 357–366.

23. Shannon, T.R., et al., A mathematical treatment of integrated Ca dynamics within the ventricular myocyte. Biophysical Journal, 2004. 87(5): p. 3351–3371.

24. Morotti, S., et al., A novel computational model of mouse myocyte electrophysiology to assess the synergy between Na+ loading and CaMKII. Journal of Physiology-London, 2014. 592(6): p. 1181–1197.

25. Sarkar, A.X., D.J. Christini, and E.A. Sobie, Exploiting mathematical models to illuminate electrophysiological variability between individuals. Journal of Physiology-London, 2012. 590(11): p. 2555–2567.

26. Britton, O.J., et al., Experimentally calibrated population of models predicts and explains intersubject variability in cardiac cellular electrophysiology. Proceedings of the National Academy of Sciences of the United States of America, 2013. 110(23): p. E2098–E2105.

27. Muszkiewicz, A., et al., Variability in cardiac electrophysiology: Using experimentally-calibrated populations of models to move beyond the single virtual physiological human paradigm. Progress in Biophysics & Molecular Biology, 2016. 120(1-3): p. 115–127.

28. Capel, R.A., et al., Two-pore Channels (TPC2s) and Nicotinic Acid Adenine Dinucleotide Phosphate (NAADP) at Lysosomal-Sarcoplasmic Reticular Junctions Contribute to Acute and Chronic β-Adrenoceptor Signaling in the Heart. J Biol Chem, 2015. 290(50): p. 30087–98.

29. Garrity, A.G., et al., The endoplasmic reticulum, not the pH gradient, drives calcium refilling of lysosomes. Elife, 2016. 5.

30. Penny, C.J., et al., A computational model of lysosome-ER Ca2+ microdomains. Journal of Cell Science, 2014. 127(13): p. 2934–2943.

31. Bayliss, R.A., Actions of NAADP and other agents in cardiac myocytes [PhD Thesis], in Department of Pharmacology. 2014, University of Oxford: uuid:8463cf89-a405-4880-9aad-6fa6ebac542d.

32. Passini, E., et al., Mechanisms of pro-arrhythmic abnormalities in ventricular repolarisation and anti-arrhythmic therapies in human hypertrophic cardiomyopathy. Journal of Molecular and Cellular Cardiology, 2016. 96: p. 72–81.

33. Zhou, X., et al., In Vivo and In Silico Investigation Into Mechanisms of Frequency Dependence of Repolarization Alternans in Human Ventricular Cardiomyocytes. Circulation Research, 2016. 118(2): p. 266–278.

34. Pitt, S.J., et al., TPC2 Is a Novel NAADP-sensitive Ca2+ Release Channel, Operating as a Dual Sensor of Luminal pH and Ca2+. Journal of Biological Chemistry, 2010. 285(45): p. 35039–35046.

35. Lloyd-Evans, E., et al., Niemann-Pick disease type C1 is a sphingosine storage disease that causes deregulation of lysosomal calcium. Nature Medicine, 2008. 14(11): p. 1247–1255.

36. Lewis, A.M., et al., beta-Adrenergic receptor signaling increases NAADP and cADPR levels in the heart. Biochemical and Biophysical Research Communications, 2012. 427(2): p. 326–329.

37. Gul, R., et al., Nicotinic Acid Adenine Dinucleotide Phosphate (NAADP) and Cyclic ADP-Ribose (cADPR) Mediate Ca2+ Signaling in Cardiac Hypertrophy Induced by beta-Adrenergic Stimulation. Plos One, 2016. 11(3).

38. Davis, L.C., et al., NAADP Activates Two-Pore Channels on T Cell Cytolytic Granules to Stimulate Exocytosis and Killing. Current Biology, 2012. 22(24): p. 2331–2337.

39. Kilpatrick, B.S., et al., Direct mobilisation of lysosomal Ca2+ triggers complex Ca2+ signals. Journal of Cell Science, 2013. 126(1): p. 60–66.

40. Menteyne, A., et al., Generation of specific Ca2+ signals from Ca2+ stores and endocytosis by differential coupling to messengers. Current Biology, 2006. 16(19): p. 1931–1937.

