## Supplemental File for "Lysosomal calcium loading promotes arrhythmias by potentiating ryanodine receptor release"

### Supplementary Material

| Parameters | | Units | Value | Ref. |
|---|---|---|---|---|
| **Volume** | Total cell ($V_{tot}$) | pL | 33 | [1] |
| | Lysosome ($V_{ls}$) | pL | 2% $V_{tot}$ | [2, 3] |
| **TPC** | Maximum open probability ($P_{Omax}$) | | 0.014 | [2, 4] |
| | Variance ($P_{Osd}^2$) | | 2.25 | this study |
| | Mean ($P_{Omean}$) | | 23 | [2] |
| | Leak rate ($j_{clc,leak}$, $j_{tpc,leak}$) | $s^{-1}$ | 1.13E-05 | this study |
| **Lysosome** | TPC flux density into junction ($j_{clc}$) | $s^{-1}$ | 9.708 | this study |
| | TPC flux density into cytosol ($j_{tpc}$) | $s^{-1}$ | 12.135 | this study |
| | [*NAADP*] for CTRL protocol | nM | 1 | this study |
| | [*NAADP*] for NAADP-AM protocol | nM | 15 | this study |
| | [*NAADP*] for ISO protocol | nM | 15 | this study |
| | Diffusion flux rate to junction ($J_{ls,j}$) | $um^3/s$ | 1.4219E-15 | this study |
| | Diffusion flux rate to cytosol ($J_{ls,i}$) | $um^3/s$ | 1.3858E-15 | this study |

**Table S1.** Baseline parameters for the lysosome compartment.

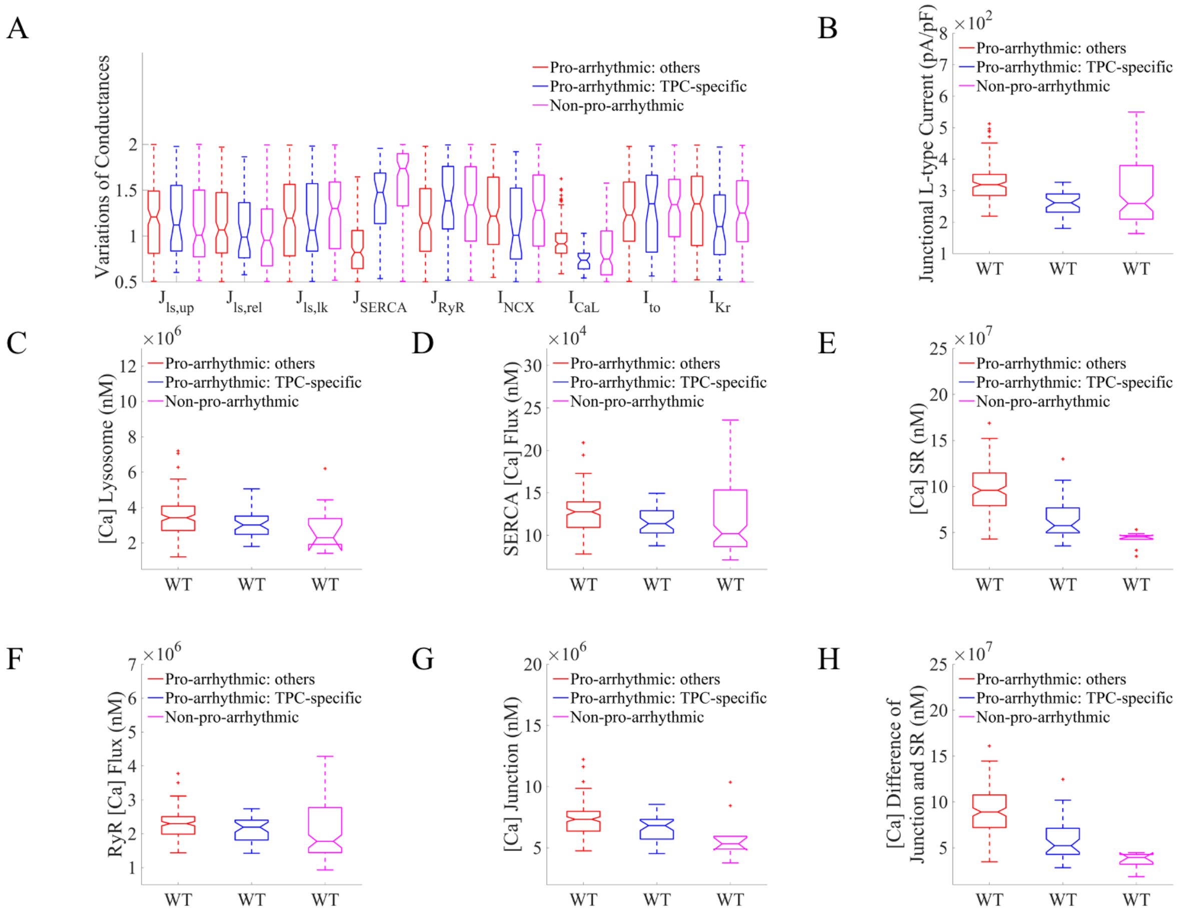


**Figure S1.** Ionic properties and calcium fluxes underlying spontaneous calcium events under hypercalcemia and sustained β-adrenergic stimulation. **S1A:** Comparison of ionic properties of TPC-specific proarrhythmic models (blue), other pro-arrhythmic models (red), and non-proarrhythmic models (magenta) in WT. **S1B-S1H:** Distributions of total (integral over entire steady-state beat) L-type calcium current, compartmental calcium concentrations, and calcium fluxes, comparing TPC-specific pro-arrhythmic models, other pro-arrhythmic models, and non-proarrhythmic models in WT.

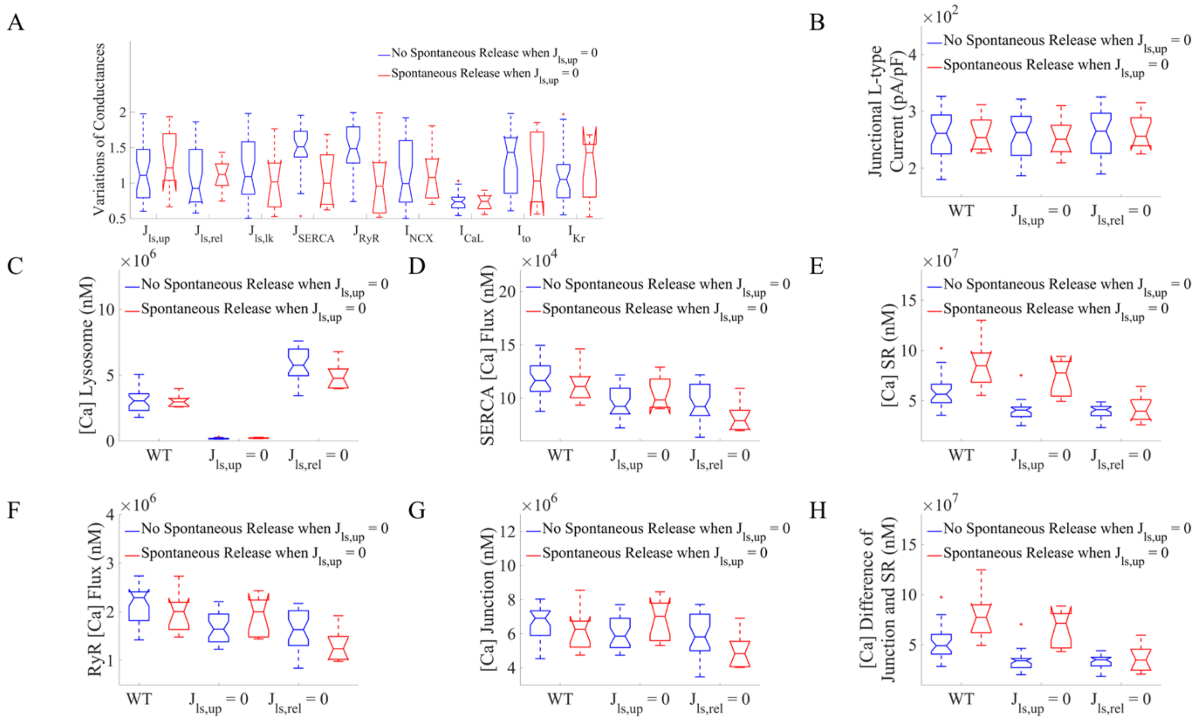


**Figure S2.** Loss of lysosomal buffer effect triggered spontaneous calcium release from SR in TPC-specific pro-arrhythmic models. **S2A**: Ionic property comparison of those either not exhibiting or exhibiting spontaneous calcium release events when blocking lysosomal calcium loading ($J_{ls,up} = 0$) in TPC-specific pro-arrhythmic models. **S2B-S2H:** Distributions of total (integral) calcium current, fluxes and concentrations over an entire beat, comparing those models either not exhibiting or exhibiting spontaneous calcium release events when blocking lysosomal calcium loading ($J_{ls,up} = 0$). Both subgroups exhibited spontaneous calcium release events in WT, while not when blocking lysosomal calcium release ($J_{ls,rel} = 0$).

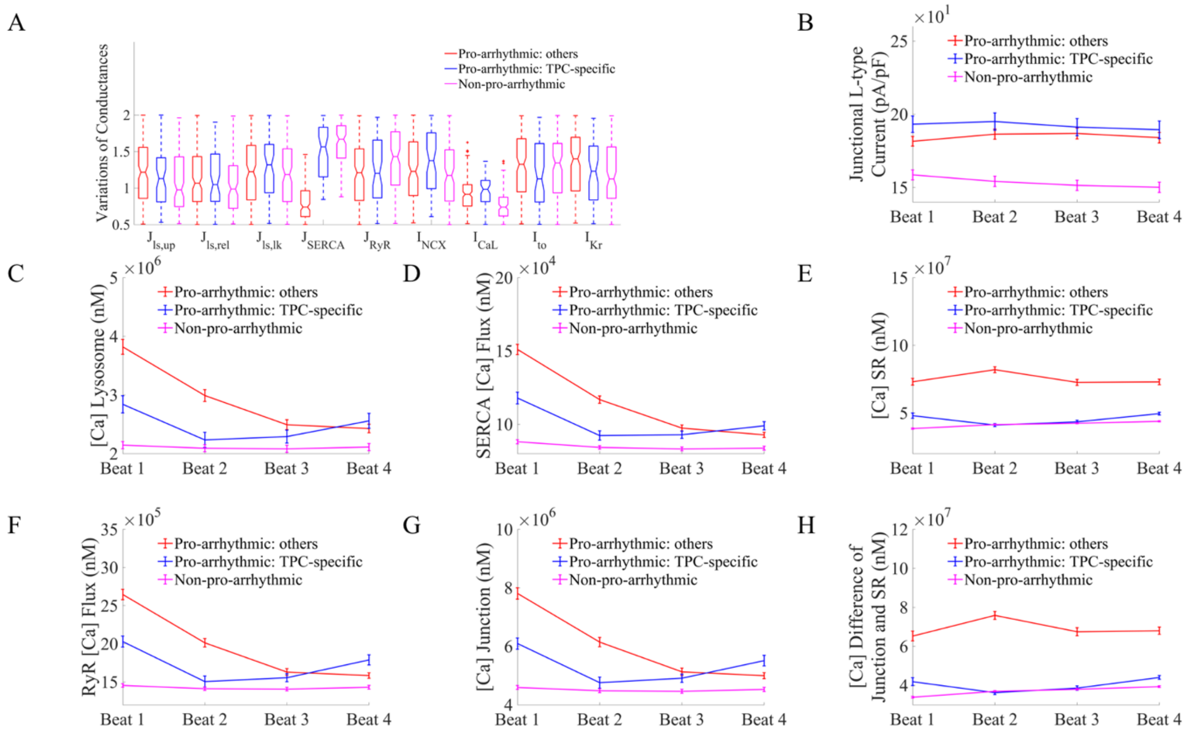


**Figure S3.** Ionic properties and calcium fluxes underlying spontaneous calcium events under fast pacing at 10 Hz and sustained β-adrenergic stimulation. **S3A:** Comparison of ionic properties of TPC-specific proarrhythmic models (blue), other pro-arrhythmic models (red), and non-proarrhythmic models (magenta) in WT. **S3B-S3H:** Distributions of total (integral over entire steady-state beat) L-type calcium current, compartmental calcium concentrations, and calcium fluxes, comparing TPC-specific pro-arrhythmic models, other proarrhythmic models, and non-proarrhythmic models in WT.

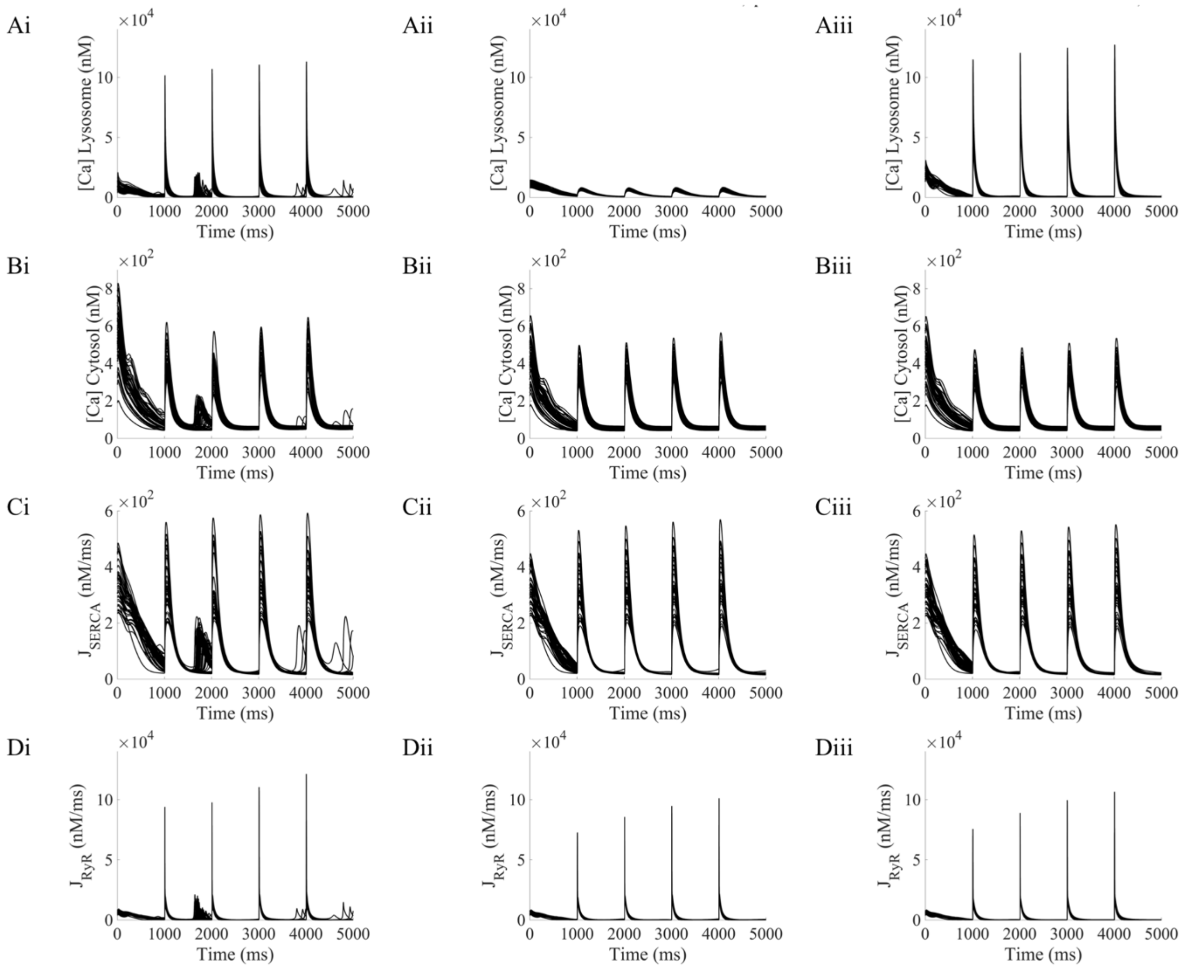


**Figure S4.** Lysosomal calcium release promotes spontaneous calcium release in TPC-specific pro-arrhythmic profiles under fast pacing and β-adrenergic stimulation, by increasing the junctional-SR calcium gradient. **S4Ai-S4Di:** Lysosomal calcium concentration, cytosolic calcium concentration, and SR reuptake and release fluxes, respectively, in basal conditions. **S4Aii-S4Dii:** Calcium concentrations and fluxes under lysosomal uptake block ($J_{ls,up} = 0$). **S4Aiii-S4Diii:** Calcium concentrations and fluxes under lysosomal release block ($J_{ls,rel} = 0$). All results presented under preliminary fast pacing at 25 Hz.

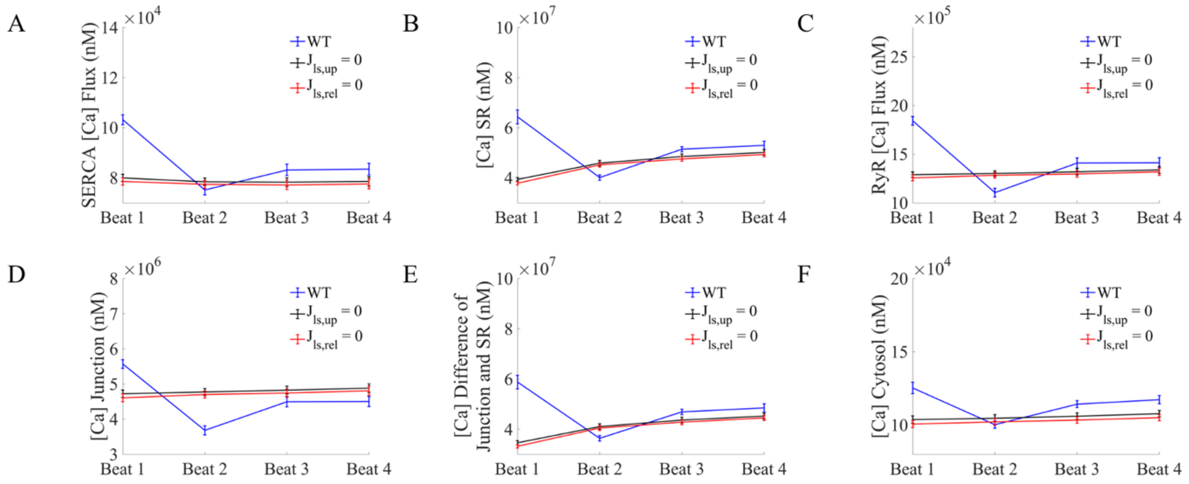


**Figure S5.** Loss of each lysosomal flux reduced spontaneous calcium release from SR in TPC-specific pro-arrhythmic models under fast pacing and β-adrenergic stimulation. **S5A-S5F:** Distributions of total (integral) calcium fluxes and concentrations per beat in the transition to slow pacing (data presented as mean±SD), comparing scenarios in WT, blocking lysosomal calcium loading ($J_{ls,up} = 0$), and blocking lysosomal calcium release ($J_{ls,rel} = 0$). Beats are numbered from beat 0 (not shown, as state is representative of fast pacing). All results presented under initial fast pacing at 25 Hz.

#### Supplemental References


1. Morotti, S., et al., *A novel computational model of mouse myocyte electrophysiology to assess the synergy between Na+ loading and CaMKII.* Journal of Physiology-London, 2014. **592**(6): p. 1181-1197.
2. Penny, C.J., et al., *A computational model of lysosome-ER Ca2+ microdomains.* Journal of Cell Science, 2014. **127**(13): p. 2934-2943.
3. Aston, D., et al., *High resolution structural evidence suggests the Sarcoplasmic Reticulum forms microdomains with Acidic Stores (lysosomes) in the heart.* Sci Rep, 2017. **7**: p. 40620.
4. Pitt, S.J., et al., *TPC2 Is a Novel NAADP-sensitive Ca2+ Release Channel, Operating as a Dual Sensor of Luminal pH and Ca2+*. Journal of Biological Chemistry, 2010. **285**(45): p. 35039-35046.
